# Arginine depletion through ADI-PEG20 to treat argininosuccinate synthase deficient ovarian cancer, including small cell carcinoma of the ovary, hypercalcemic type

**DOI:** 10.1101/845586

**Authors:** Jennifer X. Ji, Dawn R. Cochrane, Basile Tessier-Cloutier, Shary Chen, Germain Ho, Khyatiben V. Pathak, Isabel Alcazar, David Farnell, Samuel Leung, Angela Cheng, Christine Chow, Shane Colborne, Gian Luca Negri, Frieder Kommoss, Anthony N. Karnezis, Gregg B. Morin, Jessica N. McAlpine, C. Blake Gilks, Bernard E. Weissman, Jeffrey M. Trent, Lynn N. Hoang, Patrick Pirrotte, Yemin Wang, David G. Huntsman

## Abstract

**Purpose:** Many rare ovarian cancer subtypes such as small cell carcinoma of the ovary, hypercalcemic type (SCCOHT) have poor prognosis due to their aggressive nature and resistance to standard platinum and taxane based chemotherapy. The development of effective therapeutics has been hindered by the rarity of such tumors. We sought to identify targetable vulnerabilities in rare ovarian cancer subtypes.

**Experimental Design:** We compared the global proteomic landscape of six cases each of endometrioid ovarian cancer (ENOC), clear cell ovarian cancer (CCOC), and SCCOHT to the most common subtype high grade serous ovarian cancer (HGSC) to identify potential therapeutic targets. Immunohistochemistry of tissue microarrays were used as validation of ASS1 deficiency. The efficacy of arginine-depriving therapeutic ADI-PEG20 was assessed *in vitro* using cell lines and patient derived xenograft mouse models representing SCCOHT.

**Results:** Global proteomic analysis identified low ASS1 expression in ENOC, CCOC, and SCCOHT compared to HGSC. Low ASS1 levels were validated through IHC in a large patient cohort. The lowest levels of ASS1 were observed in SCCOHT, where ASS1 was absent in 2/15 cases, and expressed in less than 5% of the tumor cells in 8/15 cases. ASS1 deficient ovarian cancer cells were sensitive to ADI-PEG20 treatment regardless of subtype *in vitro*. Furthermore, in two cell line mouse xenograft models and one patient derived mouse xenograft model of SCCOHT, once a week treatment of ADI-PEG20 (30mg/kg and 15mg/kg) inhibited tumor growth *in vivo*.

**Conclusions:** Preclinical *in vitro* and *in vivo* studies identified ADI-PEG20 as a potential therapy for patients with rare ovarian cancers including SCCOHT.

**Translational relevance:** Many rare ovarian cancers lack effective management strategies and are resistant to the standard platinum- and taxane-based chemotherapy. Thus, for a rare ovarian cancer subtype like small cell carcinoma of the ovary, hypercalcemic type (SCCOHT) - an aggressive malignancy affecting young women in their twenties, effective targeted therapeutics are urgently needed. We used global proteomics to identify a deficiency in arginosuccinate synthase (ASS1) as a common feature among some rare ovarian cancer subtypes. Using *in-vitro* and *in-vivo* models, we demonstrated that the arginine-depriving investigational agent ADI-PEG20 effectively inhibited cell growth in ASS1 deficient ovarian cancers including SCCOHT, establishing it as a potential therapeutic agent for rare ovarian cancer subtypes deficient in ASS1. Further clinical investigation is warranted.

## Introduction

Ovarian cancer is the fifth most common cause of cancer death in women, accounting for more than 22,240 new diagnoses and 14,070 deaths in the United states in 2018 (1). Ovarian cancer can be categorized into epithelial cancers which mostly include high grade serous (HGSC), endometrioid (ENOC), and clear cell ovarian carcinomas (CCOC), and non-epithelial cancers that arise from ovarian germ cells, sex cord cells, stromal cells, or have unknown origin. Each subtype has unique histological and molecular characteristics with distinct cells of origin, suggesting that they are, and should be managed as, different diseases (2). Nonetheless, standard of care for all ovarian cancer subtypes, aside from surgical resection, is radiotherapy and taxane/platinum chemotherapeutics. Less common subtypes such as late-stage CCOC and small-cell carcinoma of the ovary, hypercalcemic type (SCCOHT) are often not responsive to cytotoxic chemotherapy. HGSC, the most common subtype, accounting for over 70% of all cases, has benefited from research efforts leading to the development of targeted therapeutics such as PARP inhibitors (3). However, these targeted therapeutics have limited applicability in treating patients with ovarian cancer subtypes beyond HGSC. Therefore, there is an urgent unmet need to identify novel therapeutic targets for these uncommon ovarian cancers.

While HGSC is associated with mutations in tumor suppressor *BRCA1/2* and *TP53*, both ENOC and CCOC are thought to arise from ovarian endometriosis, harboring mutations in *ARID1A, PIK3CA* and *PTEN* (4, 5). ENOC has a higher incidence of *CTNNB1* mutations and mismatch repair defects compared to CCOC (6, 7). Compared to CCOC, ENOC has a less aggressive clinical course with most patients diagnosed at early stages. CCOC accounts for about 10 - 12% of all ovarian cancer cases and is considered a high-grade malignancy. While having a better prognosis when diagnosed at early stages, about 33% of CCOC patients present at late stage, and have the worst outcome among all epithelial ovarian cancer subtypes (8). The anti-angiogenesis agent sunitinib was found to have minimal activity in refractory CCOC in a Phase II study (9). Further clinical research efforts evaluating agents inhibiting angiogenesis (NCT02866370) and overactive PI3K pathway (NCT01196429) in CCOC are underway with a recent focus on immunotherapy (10). Despite these advancements, effective and affordable targeted therapeutics for CCOC are still lacking.

SCCOHT is a highly aggressive cancer affecting young women, having a median age of diagnosis at 24 years old. The cellular of origin of SCCOHT remains unclear. The tumor is resistant to conventional chemotherapeutics and unfortunately, most patients succumb to their disease within 2 years of diagnosis (11). SCCOHT is characterized by a dual loss of SMARCA2 and SMARCA4, two mutually exclusive ATPases of the SWI/SNF chromatin remodeling complex (12). Notable recent development in therapeutics for SCCOHT include EZH2 inhibitor (13), HDAC inhibitor (14),ponatinib (15), and a CDK4/6 inhibitor (16) which show promising efficacy in preclinical models. Clinical translation of these molecules is highly anticipated. Targeted therapeutics should continue to be pursued for this highly aggressive cancer to maximize patient survival. Other non-epithelial ovarian cancers include granulosa cell tumor (GCT) and Sertoli Leydig cell tumor (SLCT). Both GCT and SLCT can have an indolent course, but at late stage and upon recurrence, the outcome is poor due to the lack of response to chemotherapy(17, 18).

To identify common vulnerabilities in rare ovarian cancers to support a basket trial design, we surveyed the global proteomic landscape of CCOC, ENOC, SCCOHT, and HGSC. We identified low/absent levels of arginosuccinate synthase (ASS1) in rare cancer subtypes including SCCOHT, a portion of CCOC, ENOC, and SLCT. In HGSC, decreased ASS1 expression has been associated with platinum resistance (19). Furthermore, Cheon *et al* reported that ASS1 protein expression by IHC overall was high in HGSC and lower in CCOC, ENOC, and mucinous cancers (20). While ASS1, the rate limiting enzyme in intracellular arginine synthesis, is expressed in most normal tissues (21), cancers low in ASS1 are auxotrophic for arginine (22). This vulnerability has been explored by using ADI-PEG20, a PEGylated arginine deiminase effectively depleting extracellular arginine (23). ADI-PEG20 is currently under Phase 1 - 3 investigation in various malignancies as a monotherapy and in combination with cytotoxic chemotherapy (22). To date, there are no clinical trials of ADI-PEG20 in ovarian cancers, possibly due to the observation that most HGSC have high ASS1 expression, but opportunities for stratification for more rare ovarian cancer subtypes remain unexplored. In this study, we showed that ADI-PEG20 is an effective therapy in preclinical models of uncommon ovarian cancer subtypes deficient in ASS1, including CCOC and SCCOHT.

## Material and Methods

### Cell lines

Cell lines representing CCOC (JHOC 5, JHOC 7, JHOC 9, OVTOKO, ES2), ENOC (IGROV1), dedifferentiated ovarian cancer (TOV112D), and SCCOHT (SCCOHT1, BIN67, COV434) were grown in RPMI with 5% FBS. HGSC cell line OVCAR3 was cultured in 199/105 medium supplemented with 5% FBS. SCCOHT1 cell line was provided by Dr. Ralf Hass (24), BIN67 were provided by Dr. Barbara Vanderhyden, and COV434 cells were provided by Dr. Mikko Anttonen. JHOC 5, JHOC 7, and JHOC 9 were obtained from the RIKEN Cell Bank. ES2 and TOV112D were obtained from ATCC. IGROV1 was obtained from the NCI cell bank whereas OVTOKO was obtained from the JCRB cell bank. Cells were maintained in an incubator with 5% CO2 at 37°C, were all STR validated and negative for mycoplasma. Early passages were used for experiments (passages 3 – 10 from thawing).

### Proteomic analyses

Global proteomic data for 6 cases each of HGSC, CCOC, and ENOC were obtained from previous publication (25). An additional six cases each of SCCOHT and HGSC were analyzed using SP3-CTP followed by the PECA bioinformatic pipeline as previously described (25). For this analysis, two 10-μM scrolls of FFPE tissue were used. The new cases were analyzed in two TMT-11 plexes. Each TMT plex contained a pooled internal standard (PIS) generated from pooling equal portions from each case included in this study at the peptide level. The PIS was used to normalized between plexes.

### Patient cohorts

Tissue microarrays (TMA) containing formalin-fixed, paraffin-embedded (FFPE) CCOC (n = 28), ENOC (n = 27), HGSC (n = 207), SCCOHT (n = 15), low grade serous (n = 9), adult granulosa cell tumor (n = 35), juvenile granulosa cell tumor (n = 8), and Sertoli-Leydig cell tumor (n = 17) were as previously described (12, 26, 27). The TMA containing endometrial endometrioid cases were included in a study by McConechy et al (6) as previously described (28). An additional 67 cases of CCOC were obtained from the OVCARE tissue bank and Vancouver General Hospital (VGH) archives, subsequently reviewed by a pathologist to confirm diagnosis (BTC, LNH). An additional 28 cases of Sertoli-Leydig cell tumor were reviewed by a pathologist (CBG). Duplicate 0.6mm cores from each case were used for tissue microarray construction using a tissue microarrayer (TMArrayer by Pathology devices). Informed patient consent was obtained under the OVCARE tissue bank protocol approved by the research ethics board (REB) (H05-60119). Use of VGH archival tissues was indicated under approved REB protocol (H02-61375).

### Immunohistochemistry and scoring

Immunohistochemistry (IHC) was performed on 4μm sections using the Ventana Discovery automated stainer (Ventana Medical Systems, Tuscon, AZ, USA) at the Genetic Pathology Evaluation Center. Staining was performed using antibodies to ASS1 (Rabbit polyclonal, Sigma, HPA020934), and Ki67 (Rabbit monoclonal SP6, Thermo, RM-9106-S0). For Ki67, anti-rabbit secondary antibody (Vector Biotin, Anti-rabbit, 1:300 with Discovery Diluent) was manually applied and incubated for 32 minutes then visualized with DAB detection kit. All TMAs were scored by pathologists (BTC, LNH). Histoscores were calculated by multiplying the average staining intensity (0 negative, 1 low, 2 moderate, and 3 intense) in tumor cells by percent tumor cells staining in 10% increments, resulting in histoscores from 0 – 300. In cases which had multiple regions represented on TMA, the highest histoscore was used. Ki67 score was determined by a pathologist (DF), and was defined as the percentage of positively staining tumor cells in the area of the most intense staining. Mitotic count was scored by a pathologist (DF) using whole H/E slides from FFPE tissue of mouse xenografts. Mitotic count per case was determined by the total mitotic figure in three high-power fields (0.237mm^2^/field), and reported as mitotic count per millimeter squared.

### Cell proliferation and IC50 assay

Cells were seeded in 96-well plates in triplicates (3000 - 5000 cells/well, depending on cell line) and allowed to attach for 24 hours. Cells were treated with 0.63μg/mL of ADI-PEG20 after attachment. Growth curves were monitored for 4 days by using an Incucyte ZOOM live cell imaging monitor (Essen BioScience, Ann Arbor, MI, USA). For IC50 assays, cells were seeded in 96 well plates in triplicates and allowed to attach for 24 hours. Cells were treated with 10 concentrations of ADI-PEG20 for 4 days. At the end of the experiment, cells were fixed in 10% methanol–10% acetic acid for 10 minutes and stained with 0.5% crystal violet in methanol for 10 minutes. The plates were dried overnight and dissolved in 10% acetic acid in water for 10 minutes and measured at 595nm in a spectrometer.

### Mouse Xenografts

Animal care was carried out following guidelines approved by the Animal Care Committee of the University of British Columbia (A17-0146). PDX-465 was passaged (p13) and injected subcutaneously as previously described (15). SCCOHT cell lines COV434 and SCCOHT1 were collected in 1X HBSS and prepared to a final volume of 200uL per mice with a 1:1 mixture with Matrigel (Corning, Cambridge, MA, USA). The final suspension was injected subcutaneously in the backs of 7-9-week-old female NRG (NOD.Rag1KO.IL2RγcKO) mice (2 × 10^6^/mouse). The mice were randomized when average tumor volume reaches 100 mm^3^ into saline control group and treatment groups. ADI-PEG20 (Polaris Pharmaceuticals) or saline control (200uL) was administered via intraperitoneal injection weekly for four weeks. Tumor volume and mouse weight were measured three times weekly. Tumor volume was calculated as length × (width)^2^ × 0.52. At study termination, tumors were collected, weighed, and fixed in 10% formalin for 48 hours before being embedded in paraffin.

### Statistical analysis

IC50 calculation was determined by the GraphPad PRISM software, all other statistical tests were performed in R. Statistical significance between 2 groups were calculated using the Student’s t-test. Statistical significance between three or more groups were calculated using ANOVA with post-hoc Tukey’s test when indicated. For proteomics, differential protein expression at the peptide level was obtained using PECA (25), p-values were FDR adjusted using the Benjamini–Hochberg procedure. Proteins with an absolute log2 fold change > 1 and adjusted p-value (p.adj) <0.05 were considered to be differentially expressed. Multigroup comparison of expression in boxplots was investigated using Kruskal Wallis test, with a post-hoc Dunn’s test with Benjamini Hochberg multiple testing correction. Statistical significance in pairwise expression comparison in box plots were calculated with a Whitney-Mann U test.

Additional methods may be found in the supplemental material and methods.

## Results

### Global proteomics identifies ASS1 as a low abundance protein in rare ovarian cancer subtypes

To identify putative therapeutic targets unique to uncommon ovarian cancers, we analyzed the global proteomic profiles of six cases each of CCOC, ENOC, and HGSC that we have previously characterized (25) and in a separate analysis, those of additional six cases each of HGSC and SCCOHT that we have recently profiled (unpublished data, manuscript in preparation) using SP3-Clinical Tissue Proteomics (SP3-CTP)(25). Differential expression analyses identified ASS1, the rate limiting enzyme in intracellular arginine synthesis (21), as having a significantly higher protein abundances in HGSC compared to ENOC and SCCOHT (log2 fold-change = 2.2 and 2.4 respectively, p.adj < 0.0001) (Fig. 1A, Fig. 1C) and a trend of having a higher expression in HGSC when compared to CCOC (log2 fold-change = 0.71, p.adj = 0.14, Fig. 1B). ASS1 peptides were identified in all analyzed samples with good coverage in both the original and additional proteomic sets (11 and 22 unique peptides, respectively), suggesting the differential expression of ASS1 was unlikely an artifact of mass spectrometry analysis. The protein expression *Z*-score indicated variable ASS1 protein abundance among six CCOC cases (Fig. 1D). In contrast, all six cases of ENOC had comparatively low ASS1 expression, and all HGSC exhibited relatively high protein abundance (Fig. 1D). In the HGSC and SCCOHT proteomic analysis, all but one case of HGSC had relatively high ASS1 abundance, and all six SCCOHT cases exhibited low ASS1 protein expression (Fig. 1E). These proteomic results suggest that while a majority of HGSC cases have high ASS1 expression, lower ASS1 expression may be a common feature among some rare ovarian cancer subtypes, including a subset of CCOC, the majority of ENOC, and all SCCOHT cases.

**Figure 1.**
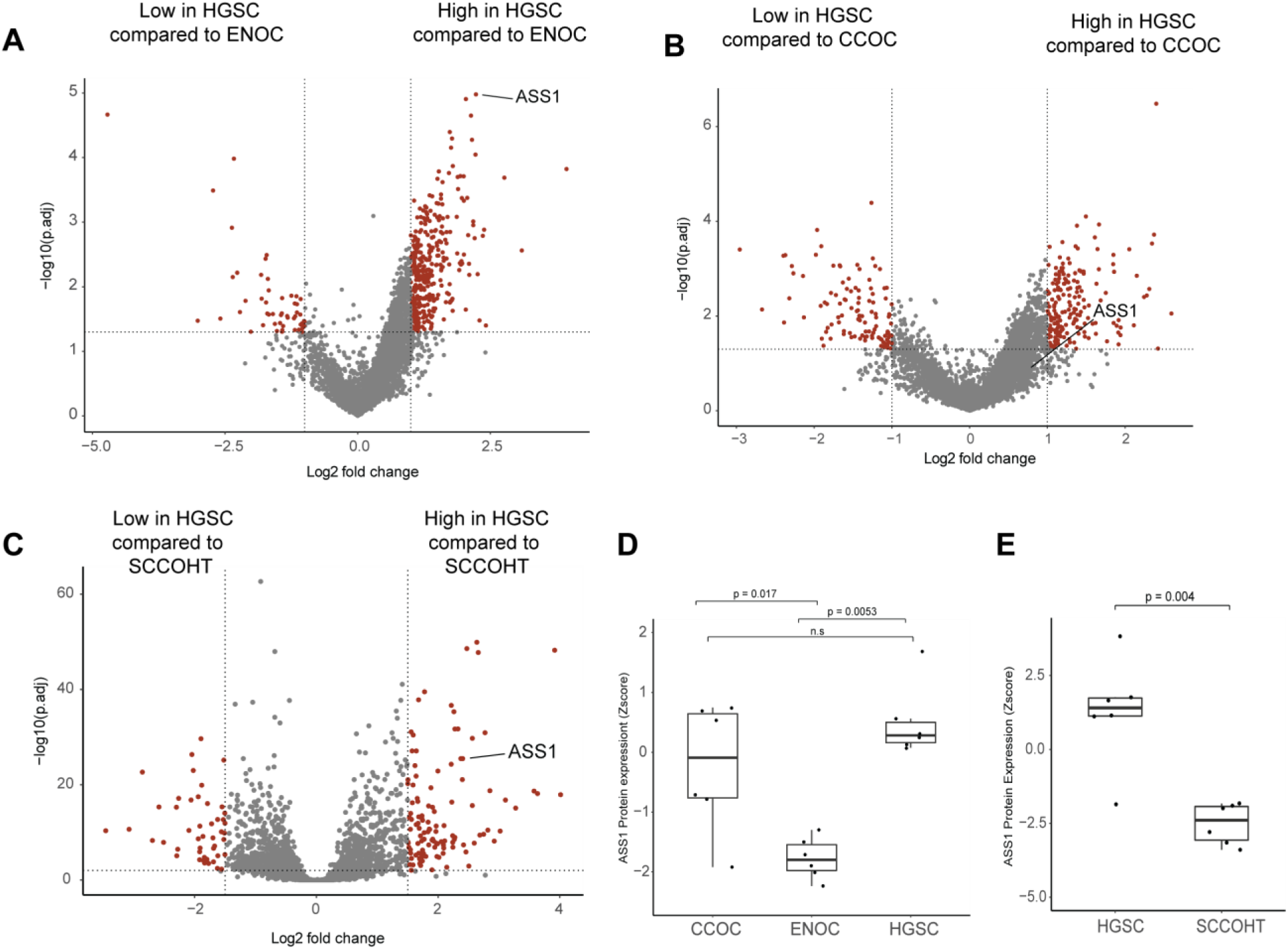
Global proteomics identifies decreased ASS1 expression in rare ovarian cancer subtypes. Proteomics data for six cases each of CCOC, ENOC and HGSC were obtained from a previous publication (25). An additional six cases each of SCCOHT and HGSC were analyzed using SP3-CTP global proteome profiling. Volcano plot showing differentially expressed proteins comparing **A**, ENOC to HGSC; **B**, CCOC to HGSC; and **C**, SCCOHT to HGSC. Significantly differentially expressed proteins are highlighted in red, and were defined as log2 fold change larger than 1 or smaller than −1, and a FDR-adjusted p < 0.05. Boxplot showing ASS1 protein expression z-score in each case of **D**, CCOC, ENOC and HGSC; and **E**, SCCOHT compared to HGSC. The statistical significance in multiple group comparisons is calculated with a Kruskal-Wallis test with a post-hoc dunn’s test with Benjamini-Hochberg correction. Pairwise comparison in figure 1E was calculated using a Mann-Whitney U test.

### Varied ASS1 expression in ovarian cancer subtypes

To validate the proteomic findings, we used IHC to survey ASS1 protein expression in an extended patient cohort including epithelial and non-epithelial ovarian cancer subtypes (Fig. 2, Table 1). ASS1 exhibited cytoplasmic staining in tumor cells, some inflammatory cells, as well as endothelial cells (Fig. 2A, Fig. 2C). In HGSC (n = 207), ASS1 was highly expressed, where 178/207 (86%) had uniformly strong expression with a histoscore of 300 (Fig. 2B). In contrast, SCCOHT had uniformly low ASS1 expression with 93% (14/15) of cases having a histoscore of less than 100 (median histoscore = 5) (Fig. 2D). Ten of fifteen cases (67%) either exhibited no ASS1 expression (n = 2), or weak staining in less than 5% of all tumor cells (n = 8). For the SCCOHT cases containing mixed histologic areas of small cell morphology, large cell morphology, and rhabdoid morphology, ASS1 staining, if present, was uniform across morphologically distinct areas, suggesting homogenous expression (Supplementary Fig. S1). In CCOC (n = 95), 69/95 (72%) expressed ASS1 strongly with a histoscore of 300 while 26/95 (28%) cases exhibited decreased ASS1 expression (histoscore <300), including 10/28 (36%) stage III/IV patients. ENOC (n = 39) had markedly lower protein expression (median histoscore = 100) (Fig. 2B). When comparing high expression (histoscore = 300) to decreased expression (histoscore <300), ASS1 expression did not correlate with overall and progression-free survival in ENOC and CCOC (Supplementary Fig. S2).

**Table 1.**
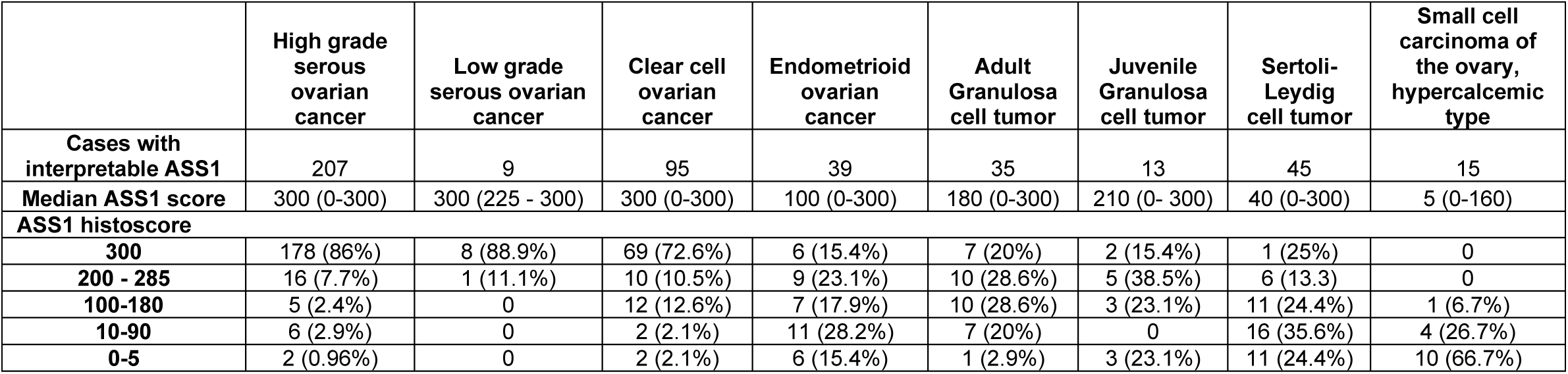
ASS1 histoscore in ovarian cancer subtypes

|  | High grade<br>serous<br>ovarian<br>cancer | Low grade<br>serous ovarian<br>cancer | Clear cell<br>ovarian<br>cancer | Endometrioid<br>ovarian<br>cancer | Adult<br>Granulosa<br>cell tumor | Juvenile<br>Granulosa<br>cell tumor | Sertoli-Leydig<br>cell tumor | Small cell<br>carcinoma of<br>the ovary,<br>hypercalcemic<br>type |
| --- | --- | --- | --- | --- | --- | --- | --- | --- |
| **Cases with**<br>**interpretable ASS1** | 207 | 9 | 95 | 39 | 35 | 13 | 45 | 15 |
| **Median ASS1 score** | 300 (0-300) | 300 (225 - 300) | 300 (0-300) | 100 (0-300) | 180 (0-300) | 210 (0- 300) | 40 (0-300) | 5 (0-160) |
| **ASS1 histoscore** |  |  |  |  |  |  |  |  |
| **300** | 178 (86%) | 8 (88.9%) | 69 (72.6%) | 6 (15.4%) | 7 (20%) | 2 (15.4%) | 1 (25%) | 0 |
| **200 - 285** | 16 (7.7%) | 1 (11.1%) | 10 (10.5%) | 9 (23.1%) | 10 (28.6%) | 5 (38.5%) | 6 (13.3) | 0 |
| **100-180** | 5 (2.4%) | 0 | 12 (12.6%) | 7 (17.9%) | 10 (28.6%) | 3 (23.1%) | 11 (24.4%) | 1 (6.7%) |
| **10-90** | 6 (2.9%) | 0 | 2 (2.1%) | 11 (28.2%) | 7 (20%) | 0 | 16 (35.6%) | 4 (26.7%) |
| **0-5** | 2 (0.96%) | 0 | 2 (2.1%) | 6 (15.4%) | 1 (2.9%) | 3 (23.1%) | 11 (24.4%) | 10 (66.7%) |

**Figure 2.**
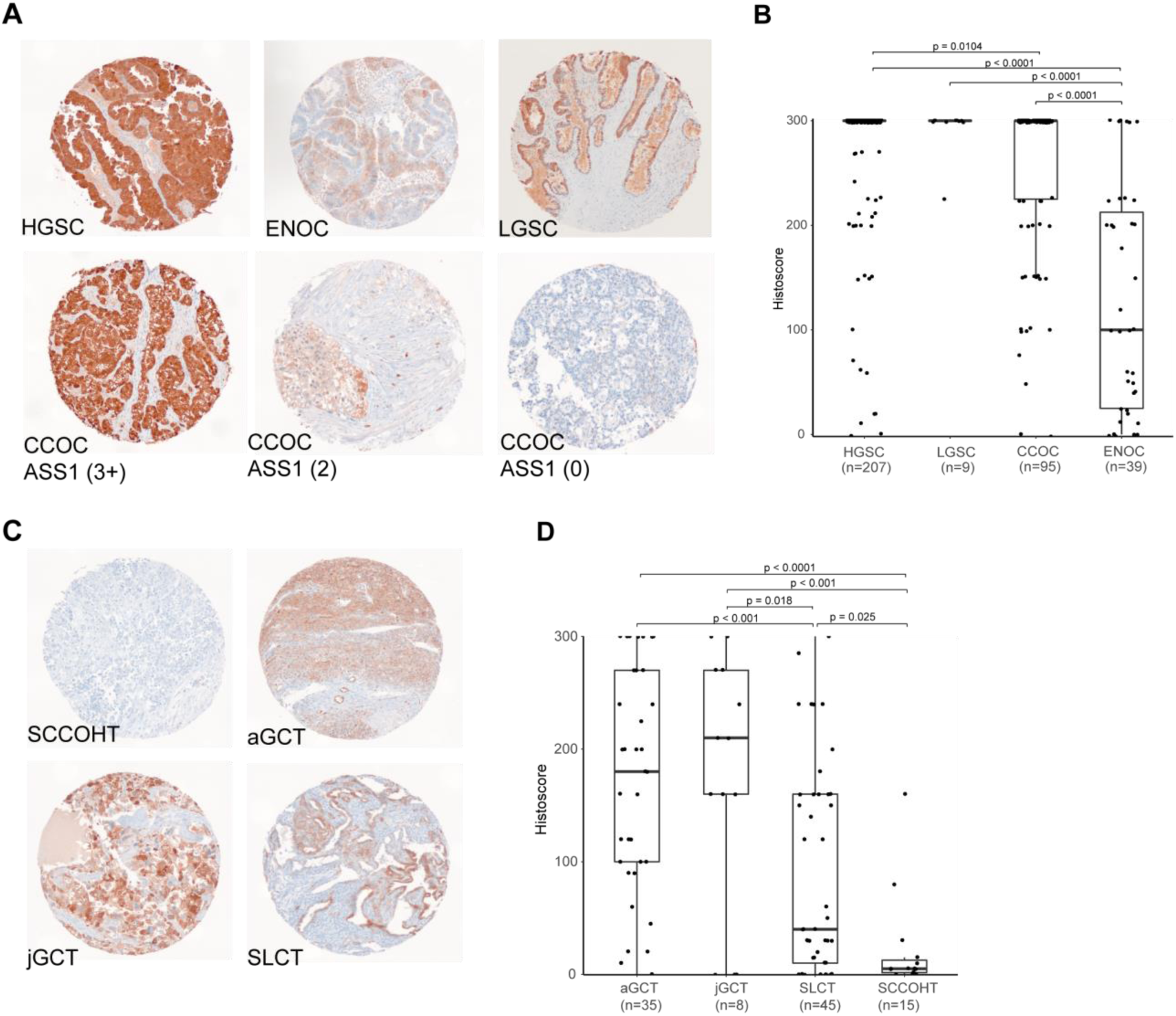
ASS1 immunohistochemistry demonstrates differential expression in ovarian cancer subtypes. Representative immunohistochemical stains in TMA cores of **A**, epithelial ovarian cancer subtypes including differential expression seen in CCOC, and **C**, non-epithelial subtypes. Corresponding boxplots depicting ASS1 histoscore distribution in **B**, epithelial subtypes and **D**, non-epithelial subtypes. Histoscore was calculated by multiplying the average staining intensity by the percentage of tumor cells staining positive. The statistical significance in multiple group comparisons is calculated with a Kruskal-Wallis test with a post-hoc dunn’s test with Benjamini-Hochberg correction.

We further determined ASS1 expression in a cohort of LGSC and sex-cord stromal cell tumors. LGSC cases (n = 9) exhibited high ASS1 expression with eight of nine cases having a histoscore of 300. In ovarian sex-cord stromal cell tumors, ASS1 expression was moderate in adult- and juvenile-granulosa cell tumors (n = 35 and 8 respectively, median histoscore = 180 and 210 respectively). ASS1 expression was decreased in Sertoli-Leydig cell tumor (n = 45, median histoscore = 40) with 60% having a histoscore <100 (Fig. 2C, 2D).

To investigate possible correlations between genomic alteration and ASS1 expression, we obtained ENOC (n=26) and CCOC (n =35) cases with available whole genome sequencing with matched RNA sequencing data (29). *ASS1* mRNA expression did not correlate with *ARID1A* or *PIK3CA* mutations in CCOC and ENOC (data not shown). All eight of 26 ENOC cases with medium to high impact *CTNNB1* mutation exhibited lower *ASS1* mRNA z-score (Supplemental Fig. S3A). We then assessed ASS1 expression using IHC in an extended local cohort of endometrioid ovarian and endometrioid endometrial cancers (EEC) for which the *CTNNB1* mutation status had been previously determined (6). Findings in the extended cohort confirmed correlation between *CTNNB1* mutation and lower ASS1 protein expression in both ENOC and EEC (*p* = 0.004 and *p* < 0.001) (Supplementary Fig. S3B). This correlation was further supported by an analysis using the EEC genomic and RNA-seq data from the Cancer Genome Atlas (*p* <0.001) (Supplementary Fig. S3C). To address whether ASS1 is silenced due to SMARCA4 inactivation in SCCOHT, we re-expressed SMARCA4 in three SCCOHT cell lines and did not observe any impact on ASS1 expression (Supplementary Fig. S4A).

### ADI-PEG20 susceptibility is specific to ASS1 deficiency in ovarian cancers

ADI-PEG20 is a PEGylated form of arginine deiminase which effectively deprives plasma arginine. It is currently being investigated in Phase 1 – 3 clinical trials targeting advanced cancers (30, 31). To assess whether ASS1-deficient ovarian cancers are sensitive to ADI-PEG20 treatment, we investigated ASS1 expression in a panel of cell lines representing a wide range of ovarian cancer subtypes. We observed ASS1 expression in the HGSC cell line OVCAR3 and differential ASS1 expression was observed in CCOC cell lines, in contrast, cell lines representing SCCOHT (BIN67, SCCOHT1, COV434), dedifferentiated ovarian cancer (TOV112D), and ENOC (IGROV1) did not express ASS1 (Fig. 3A). We then stained ASS1 on an existing TMA containing FFPE cell pellets from a panel of ovarian cancer cell lines, ASS1 IHC corresponds to ASS1 expression on western blot, supporting the utility of the antibody in an IHC capacity (Supplemental Fig. S4B). Protein expression correlated with mRNA expression in cell lines (Supplementary Fig. S4C). While *ASS1* was silenced by promoter methylation in cancers including some HGSC (19, 32), we saw promoter methylation of *ASS1* in two of three SCCOHT cell lines, but not CCOC cell lines(Supplementary Fig. S4D). Furthermore, arginine was shown to be essential for cell survival as both ASS1 expressing cell line JHOC 7 and ASS1-deficient cell line JHOC 5 exhibited growth arrest when cultured in media lacking arginine and citrulline. Citrulline, along with aspartate, is an essential precursor for the generation of arginosuccinate by ASS1 (21). The addition of citrulline rescued the growth of JHOC 7 but not JHOC 5, indicating *de-novo* arginine production is essential for ovarian cancer cell survival in arginine depleted conditions (Supplementary Fig. S4E).

**Figure 3.**
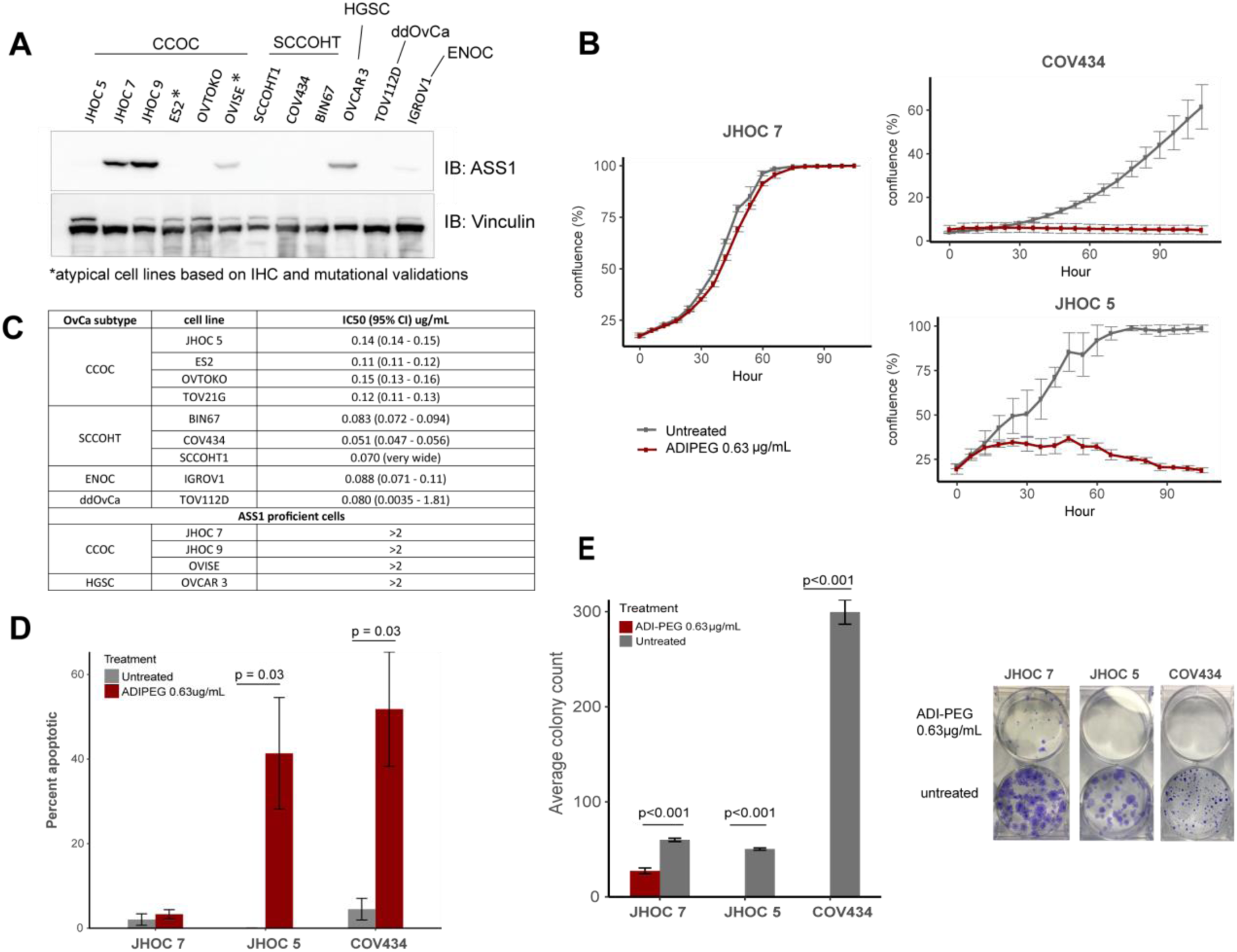
ASS1 deficient ovarian cancer are sensitive to arginine deprivation through ADI-PEG20. **A**, western blot showing differential ASS1 expression in representative ovarian cancer cell lines. **B**, cell proliferation of ASS1 negative SCCOHT cell line (COV434) and CCOC cell line (JHOC 5) compared to ASS1 expressing CCOC cell line (JHOC 7), when treated with 0.63μg/mL of ADI-PEG20; and **C**, IC50 of ASS1 deficient cell lines representing various ovarian cancer subtypes**. D**, Treatment with 0.63μg/mL of ADI-PEG20 induced apoptosis as shown by a caspase 3/7 cleavage assay in ASS1 deficient, but not in ASS1 expressing cell lines. **E**, ADI-PEG20 treatment abolished the clonogenic potential of ASS1 deficient cell lines, while decreased the clonogenic potential in ASS1-expressing cell line.

ADI-PEG20 treatment was effective in inhibiting the growth of all ASS1-deficient cell lines but not ASS1-expressing cell lines (Fig. 3B, Supplementary Fig. S5). In a 4-day assay, ASS1-deficient ovarian cancer cell lines were extremely sensitive to ADI-PEG20 regardless of subtype with the IC50 ranging from 0.051 to 0.15 μg/mL, whereas cell lines expressing ASS1 were not susceptible (Fig. 3C, Supplementary Fig. S6). To assess whether ADI-PEG20 treatment was cytotoxic, we determined the rate of cell apoptosis by measuring the activation of caspase 3/7 using a cell permeable dye that labels activated caspase3/7 followed by monitoring in a live cell monitor for three days. Two ASS1-null cell lines, JHOC 5 and COV434, exhibited a significant increase in apoptotic tumor cells (41% and 52%, respectively) compared to JHOC 7, in which no increase in apoptosis was observed (Fig. 3D). ADI-PEG20 treatment decreased colony count of the ASS1 expressing cell line JHOC 7 compared to untreated control, while completely abolishing the clonogenic potential in ASS1-deficient cells (Fig. 3E).

To evaluate the enzymatic activity of ASS1 in ovarian cancer cells, we quantified relevant metabolites in ASS1 deficient cell lines (JHOC 5 and COV434), compared to ASS1-proficient cell lines (JHOC 7 and JHOC 9) (Fig. 4A). Intracellular arginine levels were comparable between all cell lines when cultured in RPMI containing no citrulline. Upon citrulline addition, ASS1-proficient cells exhibited a significant increase in intracellular arginine levels, indicating the utilization of citrulline toward arginine generation. The immediate product of ASS1, argininosuccinate, was not present in ASS1 deficient cell lines even upon addition of citrulline, where a significant increase was seen in ASS1 positive cells in the presence of citrulline. A baseline amount of argininiosuccinate was measured in ASS1 positive cells even without the addition of citrulline, which suggests alternative mechanisms of citrulline generation, such as the nitric oxide pathways. To confirm the specificity of ADI-PEG20 sensitivity to ASS1 expression, we depleted *ASS1* by CRISPR in the ASS1-proficient cell line JHOC7 (Fig. 4B). *ASS1* knockout robustly increased the cellular sensitivity to ADI-PEG20. Furthermore, re-expression of ASS1 in JHOC5 rescued its sensitivity to ADI-PEG20 treatment (Fig. 4C), further supporting the on-target specificity of ADI-PEG20 in ovarian cancers

**Figure 4.**
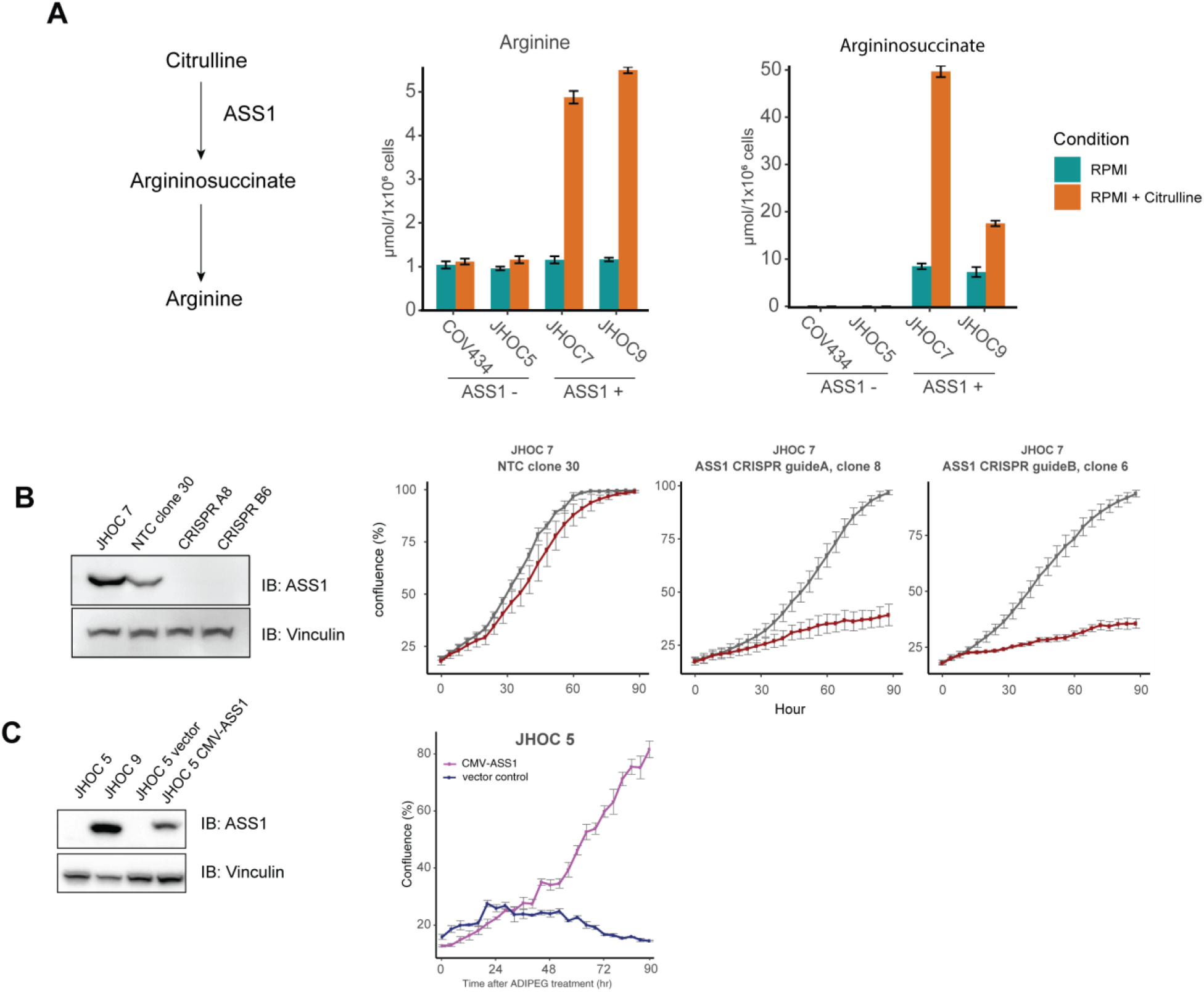
ASS1 is required for survival in arginine deplete conditions and its deficiency disrupts urea cycle function. **A**, Argininosuccinate and arginine intracellular measurements in ASS1-deficient cells (JHOC 5 and COV434), compared to ASS1-proficient cells (JHOC 7 and JHOC 9). **B**, western blot showing the knockout efficiency of two CRISPR *ASS1* knockout clones in a JHOC 7 background (A8: clone 8 using guide A, and B6: clone 6 using guide B, NTC30: empty vector negative control). and the specificity of ADI-PEG20 was confirmed in the *ASS1* knockout clones A8 and B6. **C**, Western blot showing the stable over-expression of ASS1 in JHOC5, rescuing its growth when treated with 0.63μg/mL of ADI-PEG20. All p-values were generated from student t-test. Error bars represent standard error of mean.

### SCCOHT is sensitive to ADI-PEG20 *in vivo* in cell line-derived models and patient-derived xenografts

We chose the highly aggressive SCCOHT to assess the efficacy of ADI-PEG20 *in vivo* using two subcutaneous cell line mouse model (COV434 and SCCOHT1) and one patient derived xenograft mouse model (PDX-465). Once a week treatment with ADI-PEG20 (15 mg/kg and 30 mg/kg, equivalent to about 2.5IU and 5IU respectively) (33, 34) for four weeks significantly decreased tumor growth compared to control group treated with saline (Fig. 5A). The lower dose treatment was very well tolerated. However, with the higher dose (30mg/kg), we noticed signs of toxicity including weight loss, signs of dehydration and piloerection. Some mice also exhibited enlarged kidneys and livers, but these organs showed no abnormality upon histological examination of these organs (data not shown). In the 30mg/kg treatment group for PDX-465 (n=8), we terminated one subject at day 19 due to excessive weight loss, signs of dehydration and shallow breathing. Two mice between all treatment groups had infiltrative masses above the heart which were SMARCA4 positive (data not shown), suggesting the possibility of thymic lymphomas.

**Figure 5.**
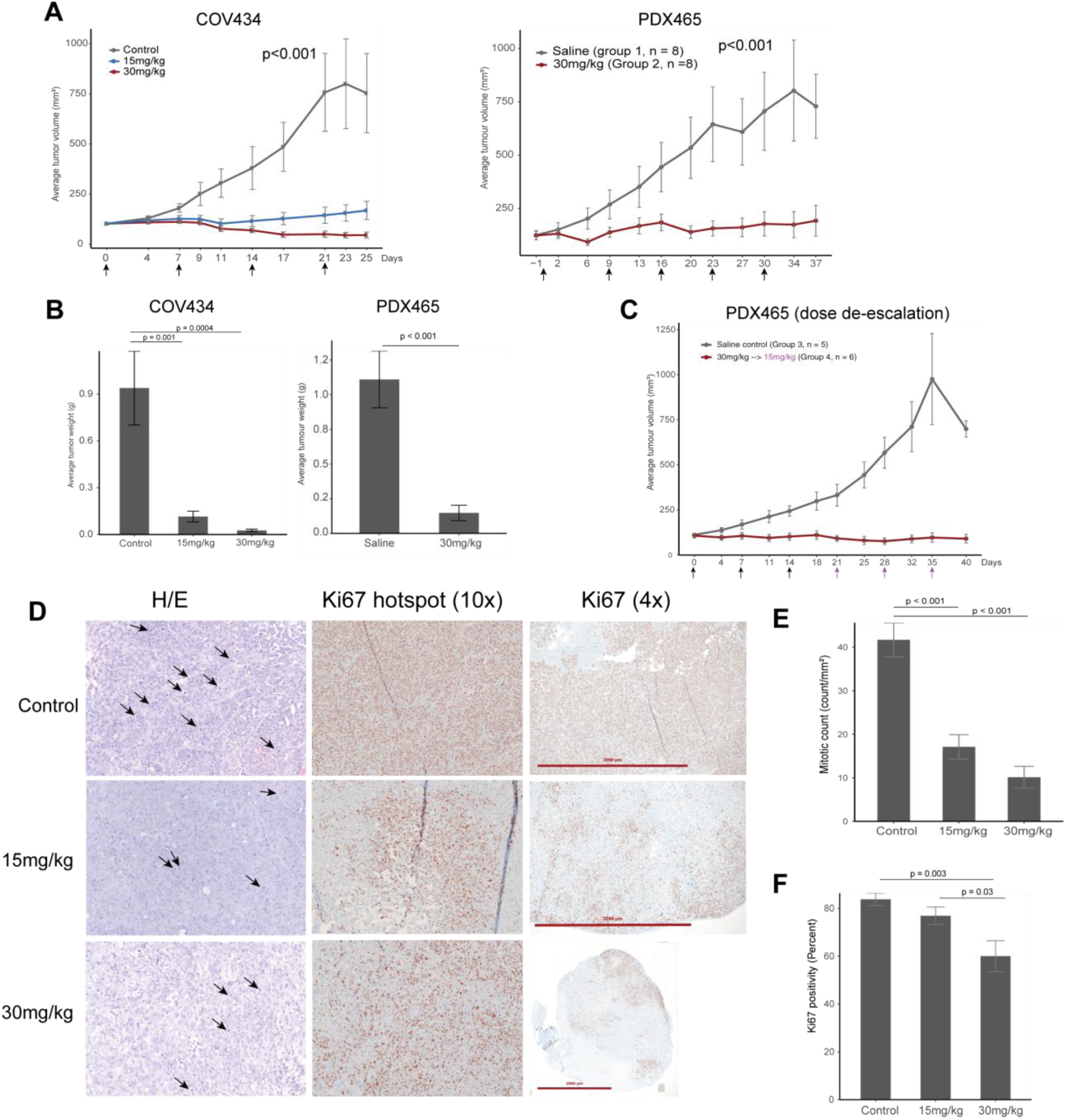
ADI-PEG20 is effective in inhibiting SCCOHT tumor growth in cell line and patient derived xenograft based *in-vivo* models. **A**, Average tumor volume of the COV434 cell line model and PDX465 patient derived xenograft model. Mice were treated once a week (indicated by arrows). **B**, average weight of tumors at study termination. **C**, average tumor volume of an additional PDX-465 experiment where subjects received three weeks of 30mg/kg (black arrows), followed by 3 weeks of 15mg/kg (magenta arrows)**. D**, histology and Ki67 immunohistochemistry of representative tumors from the control and treatment groups. Ki67 score were determined based on the area with the most intense staining. Arrows represent mitotic figures; **E**, Average mitotic count on whole sections indicating significantly dampened mitotic activity in both treatment groups. **F**, average Ki67 score between groups showing significantly decreased proliferation in the 30mg/kg group. For multiple group comparisons, significance was calculated using ANOVA, followed by a post-hoc Tukey’s test. Error bars represent standard error of mean.

In the COV434 model, tumor growth was almost completely blocked by the treatment of 15mg/kg ADI-PEG. In the 30 mg/kg group, tumors continuously decreased in size throughout four weeks (Figure 5A). By the end of the third week, three tumors in the 30 mg/kg group, and one in the 15 mg/kg group were unmeasurable using a caliper. At study termination, the average tumor weight per group in 15 mg/kg and 30 mg/kg dose were 12% and 2.7% of the control group (Fig. 5B). In the SCCOHT1 model, 30mg/kg dose significantly deterred tumor growth over a three-week treatment period (p < 0.001) (Supplemental Fig. S7A). However, the tumor weight at termination was not significant for the SCCOHT1 model (Supplemental Fig.S7A). Upon investigating ASS1 expression in the SCCOHT xenograft tumors, we noticed abundant weak to moderate ASS1 expression even in the saline control group (Supplemental Fig. S7B), contributing to the lessened effect when treated with ADI-PEG20. The identity of the cell line and xenograft was confirmed by the negativity of SMARCA4 (Supplemental Fig. S7B). The re-expression of ASS1 *in-vivo* in SCCOHT1 could reflect its increased mRNA level compared to other SCCOHT cell lines (Supplemental Fig. S4B).

Upon histological assessment of COV434 xenograft tumors, tumor morphology recapitulated SCCOHT human tumors and was similar in the treated compared to controlled groups. Mitotic count significantly decreased in both treatment groups compared to control group (p < 0.001) (Fig. 5E). Accordingly, tumors in the treated groups had foci of Ki67 staining where control group exhibited uniformly high ki67 staining (Fig. 5D). Ki67 proliferation index, scored based on the most proliferative areas, was significantly lower in 30 mg/kg group compared to both control and 15 mg/kg groups (*p* = 0.003 and *p* = 0.03 respectively). However, tumors in the treated groups appeared viable on histology. Tunnel assay also did not show increased apoptosis in the treated tumors compared to controls (Supplementary Fig. S7C). We conducted ASS1 IHC on whole sections of COV434 xenograft tumors. While no tumors expressed ASS1 in the control group, one of seven case in 30 mg/kg group and two of eight cases in the 15mg/kg group showed small foci of ASS1 re-expression (Supplemental Fig. S7D), suggesting that SCCOHT cells may regain the expression of ASS1 to develop resistance.

In concordance with our cell line model findings, ADI-PEG20 was effective in controlling tumor growth in a patient derived xenograft model – PDX-465 both with 30mg/kg, (Fig. 5A, 5B), and a dose de-escalation of 30mg/kg to 15mg/kg (Fig. 5C). This result suggests ADI-PEG20 can serve as a promising therapy for SCCOHT.

## Discussion

Effective therapy for rare and aggressive ovarian cancer subtypes is urgently needed, as they still affect thousands of women every year. Like many rare diseases, research efforts and clinical trial options for patients with rare ovarian cancers are limited. In this study, we identified ASS1 deficiency as a common vulnerability among some rare and clinically aggressive ovarian cancer subtypes, supporting the development of a rare tumor focused clinical trial using ADI-PEG20. ADI-PEG20, a pegylated form of the bacterial enzyme arginine deiminase, effectively depletes plasma arginine level (23). The agent is currently in phase 1-3 clinical trials for malignant mesothelioma, melanoma, and hepatocellular cancers, has been designated an orphan drug for malignant mesothelioma in Europe and the United States (35, 36).

The cellular origins of ovarian cancer subtypes have been much-debated; HGSC is postulated to arise from abnormal fallopian tube cells, ENOC and CCOC arise from ovarian endometriosis (37), while the cellular origin of SCCOHT remains unknown. The evolving knowledge of a diverse, extra-ovarian tissue of origin for each subtype poses a challenge in ovarian cancer research especially in comparing tumors to their corresponding normal tissues. In this study, we focus on identifying clinically actionable differences between ovarian cancer subtypes, based on molecular insights, rather than purely on cellular origins or histologic appearance. Using a validated global proteomic approach SP3-CTP (25), we compared rare ovarian cancer subtypes ENOC, CCOC, and SCCOHT to HGSC to identify decreased ASS1 expression in ENOC and CCOC compared to HGSC with confirmation through immunohistochemistry. In our patient cohort, 15% of ENOC and 72% of CCOC had intense expression of ASS1. While our ENOC results are similar to findings by Cheon *et al*, only 13% of CCOC had high levels of ASS1 in this previous report (20). This discrepancy could be due to differences in scoring metrics and antibodies used. Our results indicating high ASS1 expression in HGSC cases corroborate with previous proteomic findings in ovarian cancer cell lines (38) and immunohistochemistry findings in patient tissues (20).

In addition to epithelial ovarian cancers, we expanded our investigation into rare, non-epithelial subtypes. A global proteomics study of an additional six cases of SCCOHT and HGSC identified ASS1 as one of the most significantly differentially expressed proteins. We found universally low ASS1 expression in 15 SCCOHT cases with low heterogeneity across cores taken from different areas of the tumor In SCCOHT cell lines, we found *ASS1* silencing by promoter methylation in the COV434 and SCCOHT1 lines, but not in BIN67. However, reexpression of SMARCA4 in the cell lines did not restore ASS1 expression. Because SCCOHT possess no mutations beyond the inactivation of SMARCA4, our data suggest that the absence of ASS1 expression may represent an intrinsic feature of cell of origin of SCCOHT. We also saw low ASS1 expression in SLCT and GCT. Although both SLCT and GCT have mostly an indolent disease course, recurrent and metastatic diseases still result in poor outcome (17, 18). We did not have survival information for SLCT and GCT cases included in this study; further correlation between ASS1 expression and clinical parameters should be completed to determine whether ADI-PEG20 could be a therapy for patients diagnosed with aggressive SLCT and GCT.

The immunohistochemical survey of ovarian cancer subtypes identified possible responders to ADI-PEG20 treatment, including CCOC, ENOC, GCT, SLCT, and SCCOHT. We then showed that ASS1-deficient ovarian cancer cell lines were sensitive to ADI-PEG20 treatment regardless of subtype. ADI-PEG20 sensitivity in cell lines OVCAR3, ES2, and TOV112D agreed with previous findings (20). In addition, we report promising *in vitro* efficacy of ADI-PEG20 in a panel of CCOC and SCCOHT cell lines, for which both growth and clonogenic potential were inhibited. ADI-PEG20 treatment has previously been shown to cause cell death by apoptosis in leukemia cells (39), but could induce caspase-independent autophagy in prostate cancer cells (40). In our study, ADI-PEG20 was shown to induce ovarian cancer cell death through apoptosis.

Our observation that ASS1 expression correlated with *CTNNB1* mutation in both endometrioid ovarian cancers and endometroid endometrial cancers is curious. Because we did not find this correlation in other *CTNNB1* driven cancers such as colon adenocarcinoma (TCGA data not shown), this seems to be a gynecological cancer specific phenotype. While identifying the mechanism behind this association is beyond the scope of this study, we hope our finding can provide insight into the difficulties in clinical management associated with *CTNNB1* mutations in gynecological cancers.

In our subcutaneous COV434 mouse model, treatment with 30mg/kg (5IU) of ADI-PEG20 resulted in tumor shrinkage in all eight cases within three weeks of treatment, including one complete response. Similarly, both 30mg/kg and 30mg/kg to 15mg/kg dose deescalation in PDX-465 significantly inhibited tumor growth. While overall Ki67 staining and mitotic count decreased in the treated COV434 tumors (Fig. 5E, 5F), there were still notable Ki67 foci even in the 30mg/kg group. The *in vivo* experiments were conducted for a maximum of six weeks, most patients will receive multiple cycles of ADI-PEG20, presumably leading to sustained response. Nonetheless, the presence of residual viable tumor at the end of our study suggests that while ADI-PEG20 monotherapy induces a drastic response in SCCOHT, combination therapy may provide additional benefit in this difficult to treat disease. We observed cases with small areas of ASS1 re-expression in COV434 treatment groups, and an abundance of ASS1 expression in the SCCOHT1 model in both control and treated group, leading to therapeutic resistance. Together, these results suggest that therapeutic resistance through ASS1 re-expression could be anticipated in some SCCOHT patients where *ASS1* is silenced due to promoter methylation. In previous preclinical *in vivo* models, 5 IU of ADI-PEG20 showed efficacy in small cell lung cancer and pancreatic cancer (33, 34). A recent Phase 1/1B trial single-arm study combining ADI-PEG20 with paclitaxel and gemcitabine in 18 advanced pancreatic cancer patients showed minimal toxicity, and an overall response rate of 45% with a disease control rate of 91% (41). The encouraging clinical translation in pancreatic cancer suggests that our *in vivo* results may predict a strong response in the clinical setting for SCCOHT patients.

To date, ADI-PEG20 have been extensively studied with clinical trials in hepatocellular carcinoma, melanoma, and mesothelioma (30). ADI-PEG20 monotherapy was found to be active in a variety of cancers. Weekly treatment of ADI-PEG20 improved progression-free survival by 1.2 months in patients with refractory, chemo-resistant malignant mesothelioma (42). However, response of ADI-PEG20 monotherapy in some clinical setting have shown to be transient, related to the neutralization of the agent by host antibodies, as well as ASS1 re-expression in tumors. Instead, recent clinical trial studies focus on the combination of ADI-PEG20 with conventional chemotherapy to circumvent ADI-PEG20 resistance including studies in mesothelioma (NCT02709512), uveal melanoma (NCT02029690), and soft tissue sarcomas (NCT03449901). Early clinical activity was seen in prostate cancer and non-small cell lung cancer in a Phase 1 trial combining ADI-PEG20 with docetaxel in advanced tumors (43).

The effectiveness of ADI-PEG20 in aggressive cancers and its favorable side effect profile in drug combinations inspires confidence in its utility for aggressive and rare ovarian cancers, despite some toxicity observed in our mouse models. Recently for SCCOHT, EZH2 inhibitors and CDK4/6 inhibitors have been identified as potential therapies (13, 16). The EZH2 inhibitor Tazemetostat showed mild clinical activity in SCCOHT, with one of ten patients sustaining a partial response (44). Our results suggest that ADI-PEG20 could join the forefront of therapeutic development for SCCOHT and could be particularly useful in combination with chemotherapy or other targeted therapeutics. Combined therapy including ADI-PEG20 and immune checkpoint inhibitor in uveal melanoma is being planned (45), and a Phase 1b trial combining ADI-PEG20 and pembrolizumab in advanced solid cancers is currently recruiting (NCT03254732). In a recent case report, four SCCOHT patients were found to respond to anti-PDL1 immune check-point inhibitor (46), raising the possibility of combined ADI-PEG20 and checkpoint inhibitors for SCCOHT patients. In addition to SCCOHT, we noted low or absent ASS1 expression in 10 of 28 (36%) late-stage CCOC patients. This subset of CCOC patients may also benefit from such treatment strategies.

Furthermore, recent clinical trials combining ADI-PEG20 with conventional chemotherapy have included ASS1-positive chemotherapy refractory patients, and encouraging results have been observed in gastrointestinal cancers and pancreatic cancers regardless of ASS1 expression (41, 47). This additive efficacy has been proposed to be a result of arginine deprivation induced metabolic stress sensitizing cancers to DNA-damaging agents in an ASS1-independent manner (41). In our *in vitro* studies, ADI-PEG20 treatment significantly decreased the clonogenic potential in ASS1 expressing CCOC cell line JHOC 7 (Fig. 3). This result could indicate that combination of ADI-PEG20 and chemotherapy treatment may benefit late-stage ovarian cancer patients with or without ASS1 deficiency.

In summary, our results suggest ADI-PEG20 offers a promising therapeutic option for rare and aggressive ovarian cancer subtypes, notably in SCCOHT patients, whose clinical outcomes are otherwise dismal. Clinical evaluation through a trial focused on gynecological cancers would be an appropriate next step to determine the utility of this treatment approach.

## Author’s contributions

### Conception and design

J. Ji, D. Cochrane, Y. Wang, D. Huntsman

### Acquisition of data (Proteomics, metabolomics, *in vitro*, animal models, TMA scoring.)

J. Ji, S. Chen, G. Ho, S. Colborne, B. Tessier-Cloutier, D. Farnell, L. Hoang, K. Pathak, I. Alcazar

### Analysis and interpretation of data (e.g., statistical analysis, biostatistics, computational analysis)

J. Ji, S. Leung, G. Negri

### Writing, review, and/or revision of the manuscript

J. Ji, D. Cochrane, Y. Wang, D. Huntsman, J. McAlpine, L. Hoang, B. Tessier-Cloutier, G. Ho, G. Negri, A. Cheng, P. Pirrotte, G. Morin, B. Weissman, K. Pathak

### Administrative, technical, or material support (i.e., provided patient cases for TMA build, IHC staining)

A. Cheng, C. Chow, A. Karnezis, F. Kommoss, J. Mcalpine, CB. Gilks, G. Morin, P. Pirrotte, B. Weissman, J. Trent

### Study supervision

D. Huntsman

## Acknowledgements

We would like to thank Polaris Pharmaceuticals for providing ADI-PEG20, and OVCARE tumor bank for providing patient tissues. The authors would like to thank Derek Wong, Winnie Yang, Janine Senz, and Amy Lum for their technical support. We would like to thank Michelle Woo for assisting with REB protocols for this study.

