## Supplemental methods for "Arginine depletion through ADI-PEG20 to treat argininosuccinate synthase deficient ovarian cancer, including small cell carcinoma of the ovary, hypercalcemic type"

**Supplemental Material and Methods**

**CRISPR ASS1 Knockout**

CRISPR gRNA constructs were designed to maximize on-site specificity and minimize off-target activity using publicly available online CRISPR design algorithms. Two gRNAs were chosen, targeting Exon 5 and Exon 3 of *ASS1* transcript reference sequence. (gRNA A: 5' CACCTTTCCTGTGGCGCCGT; gRNA B: 5' ATAGCGTCTGGCGGGAGTCG). The gRNAs were cloned into the LCV2-DRX2 CRISPR backbone (Zhang Lab, MIT) and selected by ampicillin resistance in Stbl3 bacteria according to protocols developed by the Zhang Lab. DNA sequencing of the gRNA region of the LCV2-DRX2 CRISPR construct was used to verify proper gRNA ligation. Lentiviral production was performed in HEK293T cells using the PolyJET transfection agent (SignaGen, SL100688). JHOC 7 cells were seeded in 6-well plates overnight and transduced with lentivirus for 24 hours at 37°C in serum-free media with polybrene. Two gRNAs for *ASS1* and one non-targeting control gRNA transduction were performed simultaneously in different wells. Cells were expanded in 10 cm dishes and FACS sorted using the ARIA Fusion system for Ds-Red reporter expression so that single cells occupied individual wells of a 96-well plate. These cells were clonally expanded. Clones were screened using western blot to confirm knockout efficiency.

**ASS1 over-expression**

Human *ASS1*-CMV-GFP-Puro lentiviral expression vector (LV707909) and matching blank control vector (LV590) were purchased through ABM (Vancouver, Canada). Lentiviral packaging was performed with PAX2 and VSV.G viral plasmids using Trans-LT1 transfection reagent (Mirus) in early passage HEK293T cells. Transduction with *ASS1* expression and negative control vectors were performed on freshly seeded JHOC5 cells at 70% confluency. Cells were incubated with viral media containing fresh lentivirus, RPMI 1640 with 5% FBS, and 4 ug/mL polybrene for 20 hours. Cells were washed thoroughly, passaged and re-plated into fresh media (RPMI 1640 with 5% FBS) supplemented with (1ug/mL) puromycin for selection.

**Quantitative measurement of urea cycle metabolites**

Triplicate biological replicates of JHOC 5, JHOC 7, JHOC 9 and COV434 were seeded in 10 cm dishes ($9x10^5$ cells per 10cm dish) and allowed to adhere for 24 hours. Culture media was then replaced to either standard RPMI media or RPMI with 20µg/mL citrulline. Cells were grown to 70-80% confluence (approximately 3 days) and 1 million cells were collected. For metabolite measurements, cells were scraped and washed two times with cold PBS. Cell pellets were then snap-frozen and stored at -80°C until analysis. Metabolite extraction was performed using a solvent mixture of acetonitrile, methanol and water (40:40:20, v/v/v) and spiking with 10 µL of 1µM isotopically labeled internal standard mix. Each extract was incubated at -20°C for 15 min followed by centrifuging at 13,500 rpm, 4°C for 15 min. The resulting supernatant was kept at 4°C, while the remaining cell debris were subjected to secondary metabolite extraction and incubated at 4°C for 15 min, followed by centrifugation at 13,500 rpm, 4°C for 15 min. Both supernatants were combined and dried. Metabolite extracts were resupended in 100µL of water for LC-MS/MS analysis.

**LC-MS/MS analysis of urea cycle**

Urea cycle metabolites arginine and argininosuccinic acid used for method development and calibration curves were procured from Sigma Aldrich (St. Luis, USA). Isotopically labeled internal standards (IS) of the urea cycle metabolites (13C6 arginine) were obtained from Toronto chemicals (North York, Canada). Quantitative analysis of urea cycle components was performed using the stable isotope dilution method in combination with an LC-MS/MS based scheduled multi-reaction monitoring (MRM) approach on a Waters Acquity UPLC coupled to a Waters Xevo TQ-s mass spectrometer (Milford, USA). MassLynx v4.1 was employed for data acquisition. Chromatographic separation (5 µL injections) was achieved on a Synergy Hydro-RP column (100 mm x 2 mm, 2.5 µM particle size, Phenomenex, Torrance, USA) running at a column temperature of 35°C. Solvent A and B were defined as follows: A: water, 0.1% formic acid, and B: acetonitrile, 0.1% formic acid. A 9 min gradient of 100% solvent A for 0-5 min, 100% to 1% solvent A for 5-9 min at flow rate of 0.2 mL/min was used for separation. The final 5.5 min were employed for washing and equilibration. For mass spectrometry analysis, electrospray ionization was used in positive mode with a capillary voltage at 3kV, a source temperature at 150°C, and a desolvation temperature at 600 °C. The MRM transitions are detailed in the table below. Automated peak integration and quantitation were performed using TargetLynx v4.1. Metabolites were quantified (0-500 µmoles, R2=0.9) using the area-under-the-curve ratio of metabolite to internal standard. All the peaks were visually inspected. The metabolite measurements were normalized to cell counts and represented in µmoles/1x$10^6$ cells.

MRM parameters for urea cycle metabolites

| Metabolite | Precursor Adduct | Precursor m/z | Cone Voltage (V) | Collision Energy (V) | Product m/z |
|---|---|---|---|---|---|
| Arginine | M+H | 175.1 | 6 | 14,24,14 | 60.1,70.1,116.1 |
| Arginine_IS | M+H | 181.1 | 6 | 14,24,14 | 61.1,74.1,121.1 |
| Argininosuccinic acid | M+H | 291.1 | 6 | 28,18,14 | 70.2,116.1,158.4 |

**Caspase 3/7 Assay**

Cells were seeded in 96-well plates and allowed to attach for 24 hours. After attachment, cells were treated with either 0.63µg/mL of ADI-PEG with1:1000 dilution of caspase 3/7 cleavage reagent (Essen Biosciences, IncuCyte® Caspase-3/7 Green Reagent for Apoptosis), or 1:1000 dilution of DMSO as control. The cells were monitored using an IncuCyte imager for three days. Green fluorescence represented apoptotic cells, where phase data was used for confluency. Percent apoptosis was calculated as the average green area divided by the average phase area.

**Clonogenic Assay**

300 cells were seeded per 6-well plate in triplicates. The cells were allowed to attach for 24 hours prior to treatment with 0.63ug/mL ADI-PEG20. Cells were grown until visible individual colonies establish, about 14 days. Wells were fixed in 10% methanol–10% acetic acid for 10 minutes and then stained with 0.5% crystal violet in methanol for 10 minutes. The plates were

washed in water and allowed to dry overnight. Colonies were counted and the average were taken for the triplicates.

**TCGA analysis**

*ASS1* mRNA raw count and *CTNNB1* mutation status of the TCGA uterine corpus endometrial carcinoma (UCEC) (1) was downloaded from Firebrowse (http://firebrowse.org/?cohort=UCEC). The statistical significance in *ASS1* expression between *CTNNB1* mutated and wildtype cases were calculated using Wilcoxin rank sum test.

**Methylation PCR**

Genomic DNA was extracted from cell lines using the DNeasy Blood & Tissue Kit (Qiagen) and quantified by a Qubit Fluorometer (ThermoFisher). 600 ng of DNA were used in the bisulfite conversion using the EpiTect Fast Bisulfite conversion kit (Qiagen). Normal female genomic DNA (Promega, G1521) was used as negative control, and CpG methylated human genome DNA (ThermoFisher, SD1131) was used as positive control. *ASS1* methylated and unmethylated CpG primers were ordered from IDT according to a previous publication (2), where methylated sequences (M) were: forward 5′-GTAGGAGGGAAGGGG-TTTTC-3′ reverse 5′-GCAAAAAACAAATAACCCGAA-3′ and unmethylated sequences (U) were: forward 5′-GTAGGAGGGAAGGGGTTTTT-3′, reverse 5′-ACAAAAAACAAA-TAACCCAAA-3’. 2uL of Epitect-converted DNA was used in the PCR reaction. PCR was conducted on an Alpha Cycler 4 PCRmax machine (Cole-Parmer) using platinum Tag enzyme (Invitrogen). For methylated primers PCR conditions were as follows: 95$^{o}$C for 5 minutes, 45 cycles of: 95$^{o}$C for 30 seconds, 61$^{o}$C for 30 seconds, followed by 72$^{o}$C for 30 seconds. The final extensions were incubated at 72$^{o}$C for 5 mins. For unmethylated primers, the extension was held at 52$^{o}$C for 45 cycles. PCR products were run on a 2% agarose gel containing GelStar$^{TM}$ Nucleic Acid Gel Stain (Lonza, Catalog #: 50535) to visualize bands.

**Immunoblotting**

Whole-cell extracts were obtained from cell lines using RIPA buffer containing proteasome inhibitors. Antibodies against ASS1 (Sigma, HPA020934, 1:500 in 3% BSA) or vinculin (clone hVIN-1, V9131; Sigma, 1: 5000 in 3% BSA) was incubated overnight at 4$^{o}$C.

**qPCR**

Total RNA was extracted using the RNeasy kit (Qiagen) and quantified by Nanodrop. 1.25µg of total RNA was used for cDNA conversion with Superscript IV (Invitrogen). qPCR amplification was completed using Power CYBR Green master mix (Life Technology). The ASS1 primers are as previously described (3).Forward 5'-GAAGTGCGCAAAATCAAACA-3';
Reverse 5'-ATGTACACCTGGCCCTTGAG-3'

**Tunnel assay**

Tunnel assay was performed as indicated by the manufacturer using the Tunnel Assay DAB-HRP kit (Abcam, ab206386). The final DAB development was modified to five minutes, and counterstain with methylene blue was allowed to incubate for one minute.
