## Supplemental Figures 1-7 for "Arginine depletion through ADI-PEG20 to treat argininosuccinate synthase deficient ovarian cancer, including small cell carcinoma of the ovary, hypercalcemic type"

**Supplemental figures and legends**

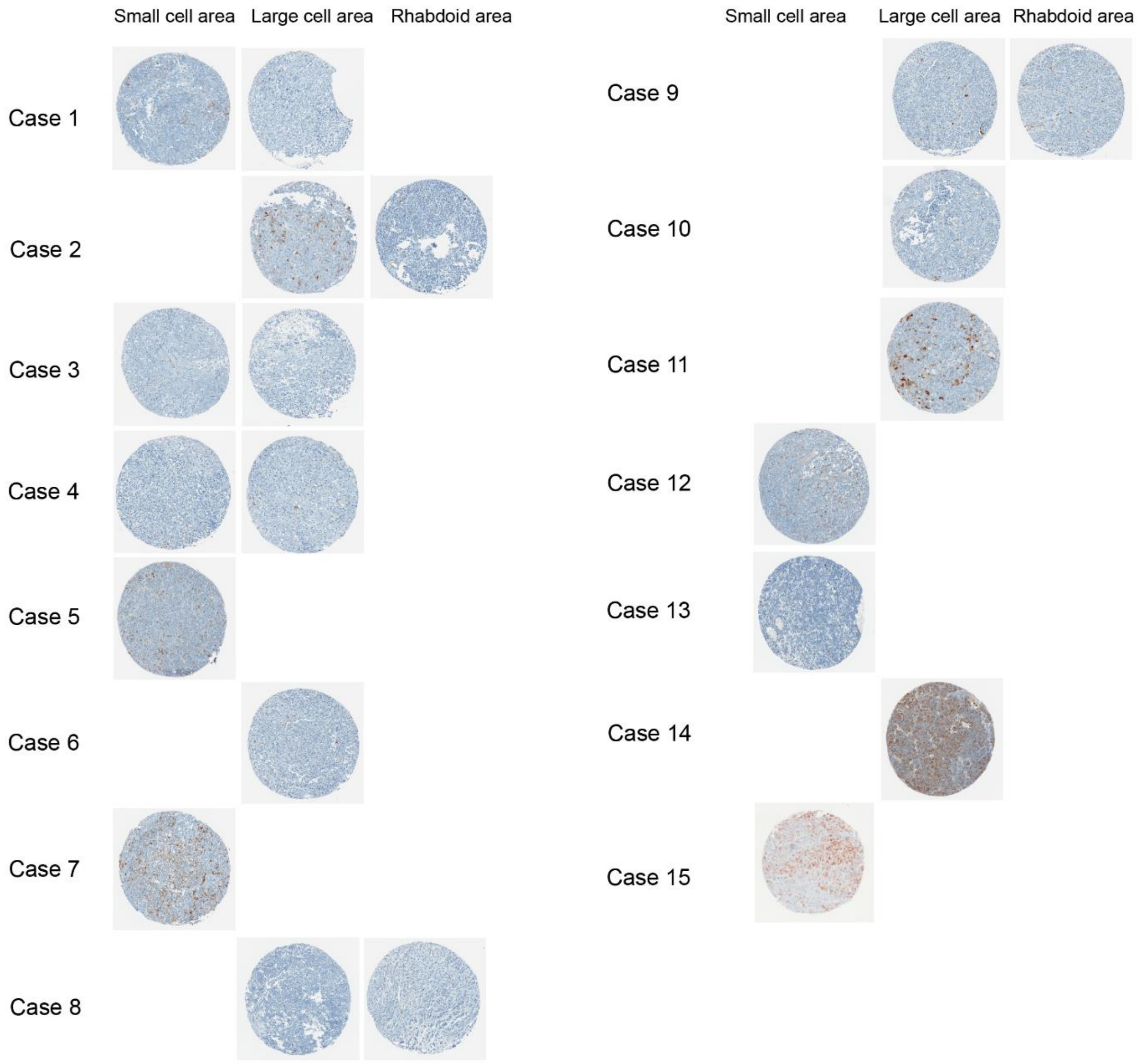


**Figure S1**. ASS1 immunohistochemistry in all 15 cases of SCCOHT. Different areas (small cell, large cell, rhabdoid) are represented for some cases. Representative cores for each area for each case are shown. Final histoscore for each case is the highest score among all areas.

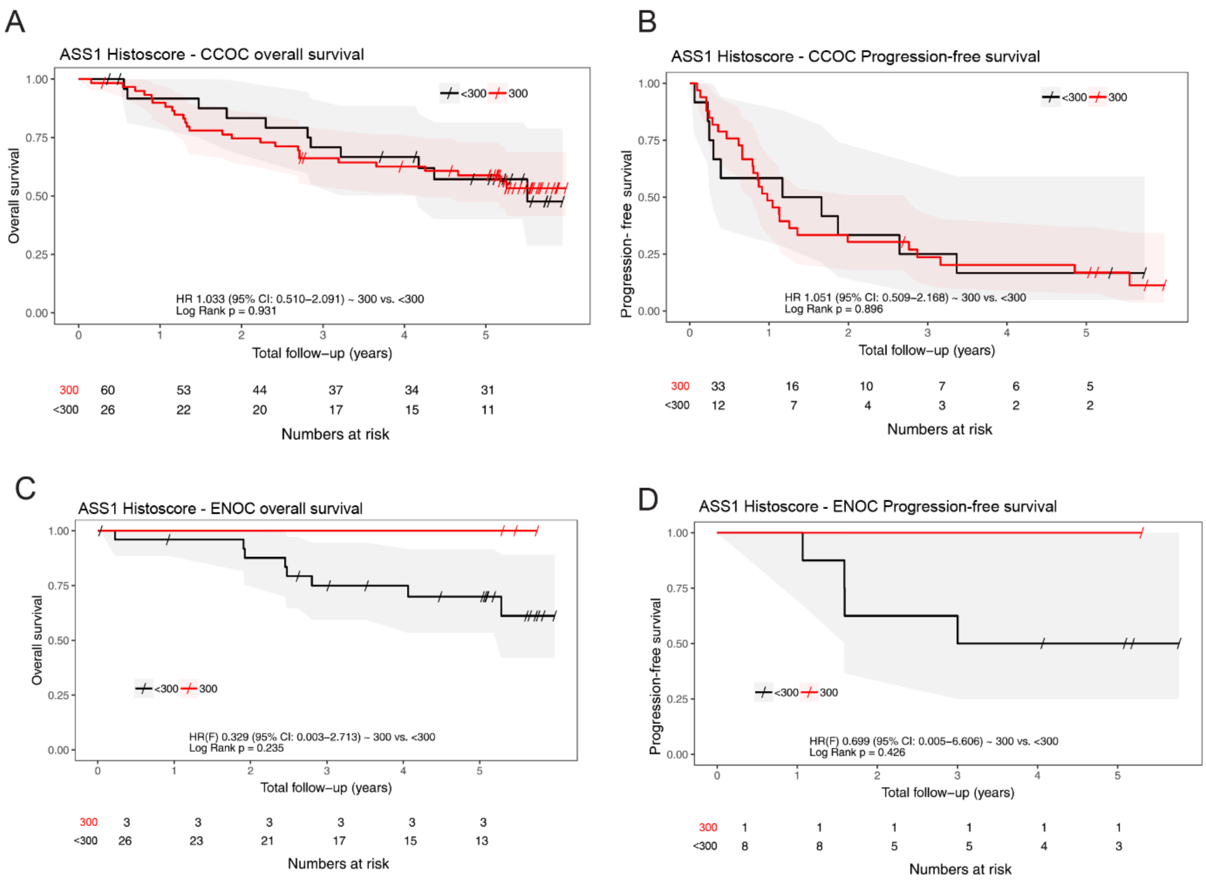


**Figure S2.** ASS1 expression does not correlate with survival in CCOC and ENOC. Kaplan-Meier plots with risk table comparing intense ASS1 expression (histoscore = 300, red line) to low/moderate ASS1 expression (histoscore <300, black line) for **A,** overall survival in CCOC**; B,** progression free survival in CCOC; **C,** overall survival in ENOC; and **D,** progression-free survival in ENOC.

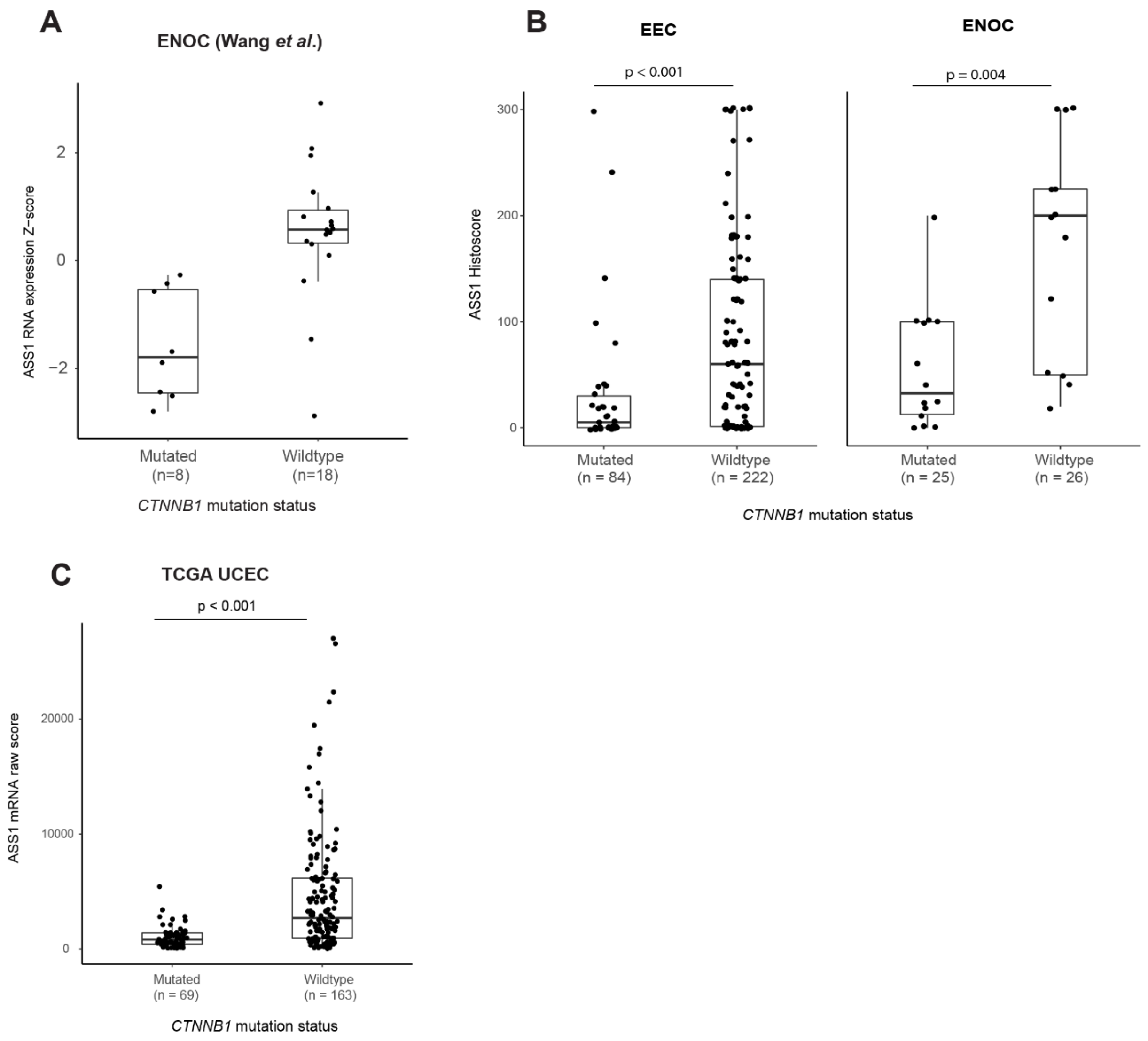


**Figure S3**. Decreased ASS1 expression correlates with *CTNNB1* mutation in ENOC and EEC. **A,** ASS1 mRNA Z-score in 26 cases of ENOC cases described in publication by Wang *et al* (29). **B,** ASS1 histoscore distribution in *CTNNB1* mutated and wildtype endometroid endometrial and endometrioid ovarian cancers. **C,** ASS1 mRNA raw counts in the TCGA endometrial cohort as related to *CTNNB1* mutation status. P values were calculated using a Wilcoxin rank sum test.

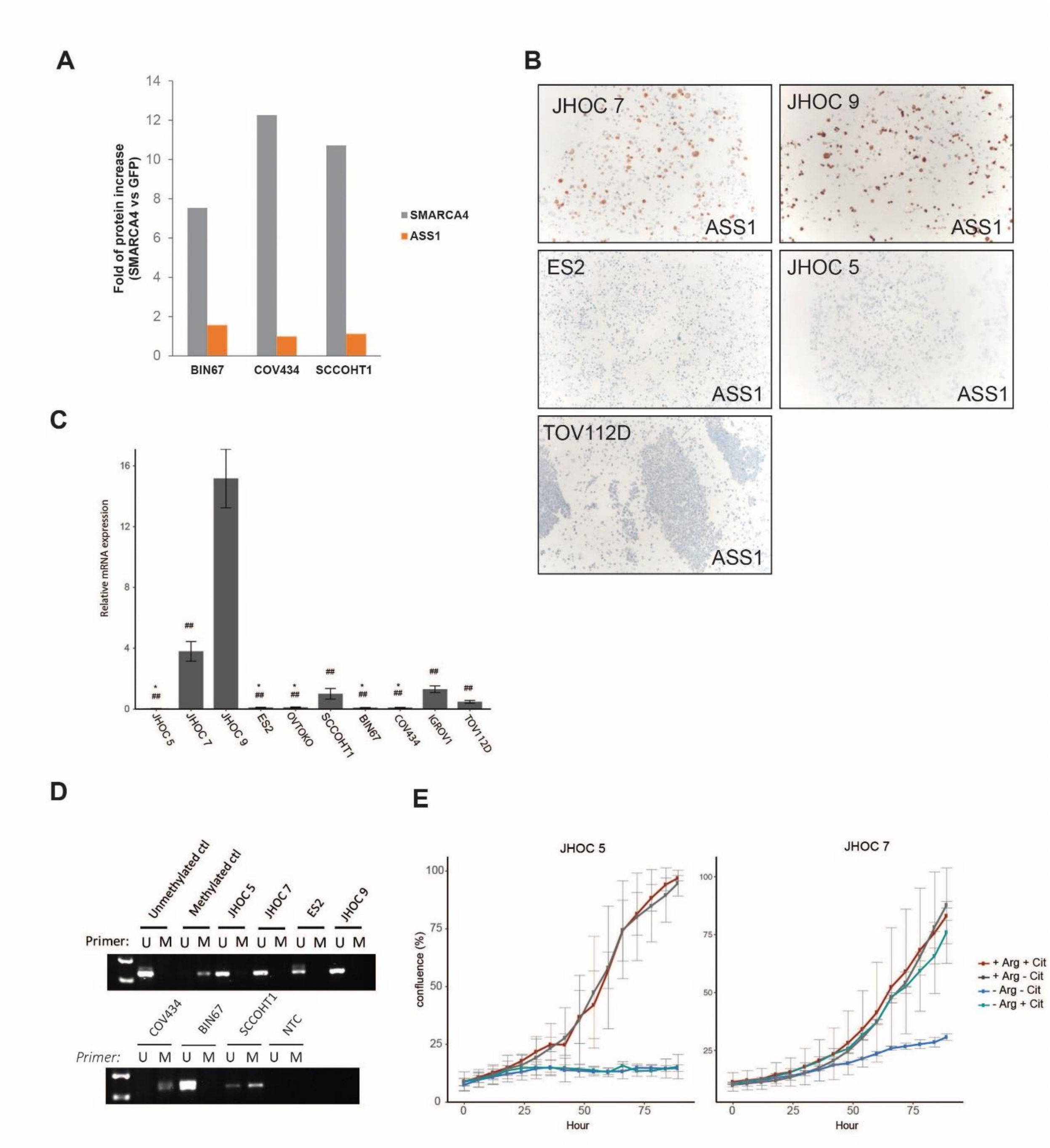


**Figure S4.** ASS1 in ovarian cancer cell lines. **A,** Re-expression of SMARCA4 does not affect the expression of ASS1. SCCOHT cells were infected with Lenti-GFP or Lenti-SMARCA4 for 24 hours followed by puromycin selection for 72 hours. Cells were then harvested for protein quantification using mass spectrometry. **B,** ASS1 immunohistochemistry on FFPE cell line pellets showing positivity in JHOC 7 and JHOC 9, and lack of expression in ES2, JHOC 5, and TOV112D. **C,** qPCR showing differential ASS1 mRNA expression levels between ovarian cancer cell lines. Difference of relative mRNA expression between cell lines is determined by one-way anova with post-hoc Tukey test. * $p<0.05$ when compared to JHOC 7; ## $p <0.001$ when compared to JHOC 9. Error bars represent standard error of mean. **D,** ASS1 promoter methylation status, all samples were run on the same gel. **E,** Cell lines are sensitive to depletion of arginine and citrulline from growth media. Addition of citrulline rescued the growth of ASS1 expressing JHOC7, but not the ASS1 deficient JHOC 5 and COV434.

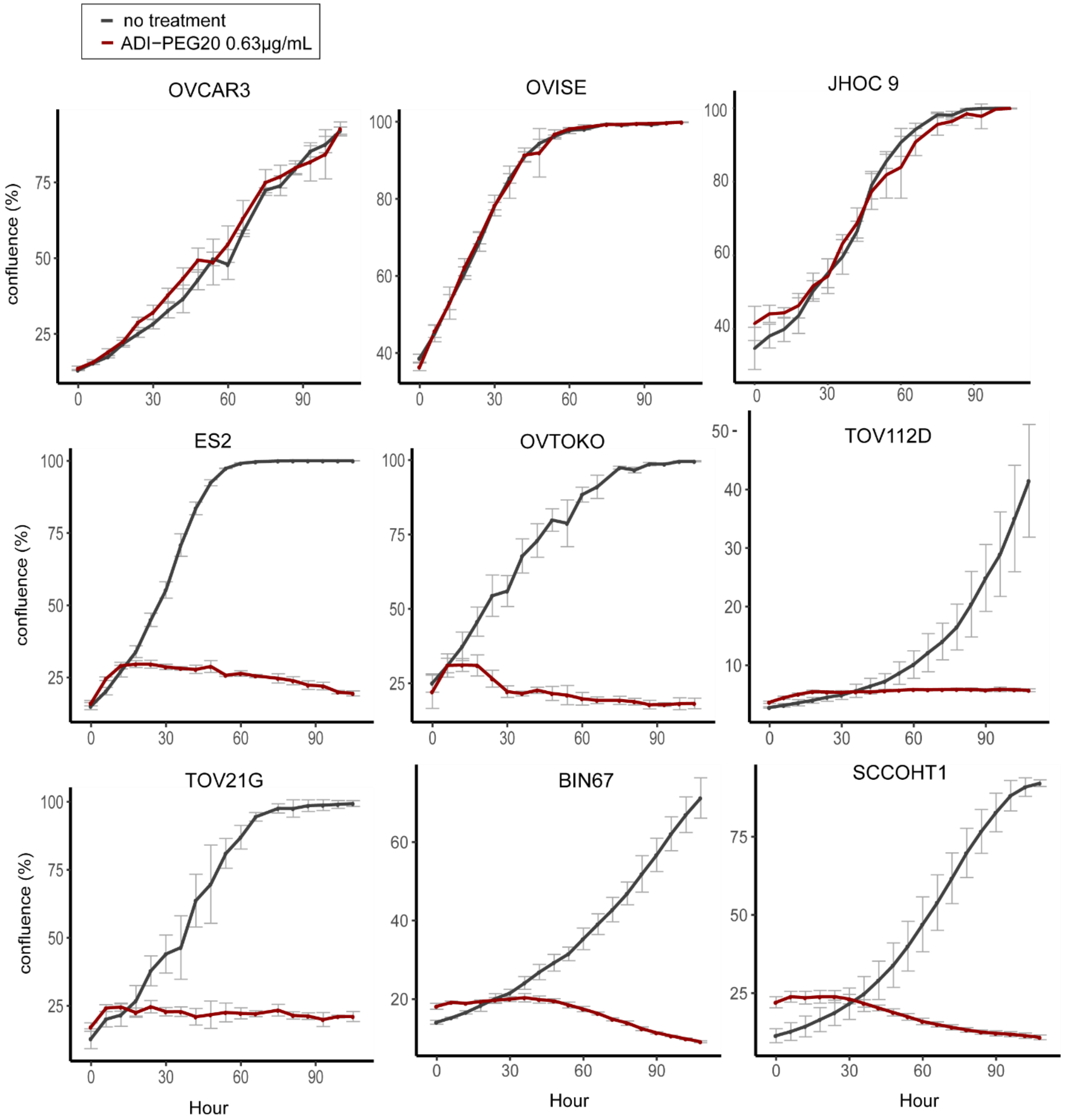


**Figure S5.** ADI-PEG20 efficacy in an extended panel of ovarian cancer cell lines. Cells were treated with 0.63 µg/mL ADI-PEG20. First row represents ASS1 expressing cell lines, whereas the rest of the cells are ASS1 deficient, therefore sensitive to ADI-PEG20 treatment. OVCAR3: HGSC cell line; JHOC 9, OVTOKO, TOV21G: CCOC cell lines. BIN67, SCCOHT1: SCCOHT cell lines; OVISE and ES2 represent atypical CCOC/ENOC based on molecular characterizations; TOV112D: de-differentiated ovarian cancer cell line. Error bars represent standard error of mean.

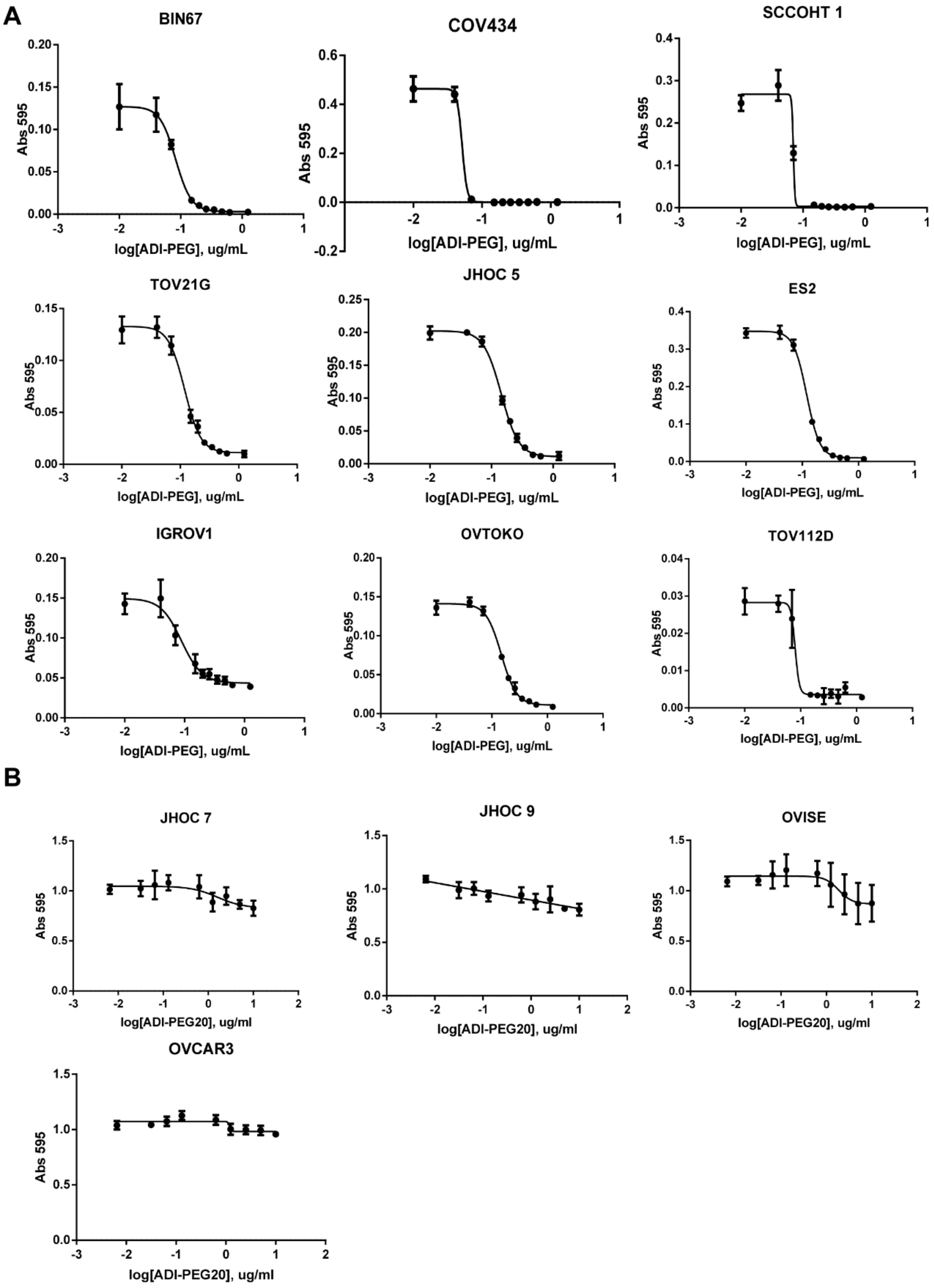


**Figure S6.** IC50 curves for ADI-PEG20 in ovarian cancer cell lines. **A,** ASS1-deficient ovarian cancer cell lines and **B,** ASS1-proficient ovarian cancer cell lines.

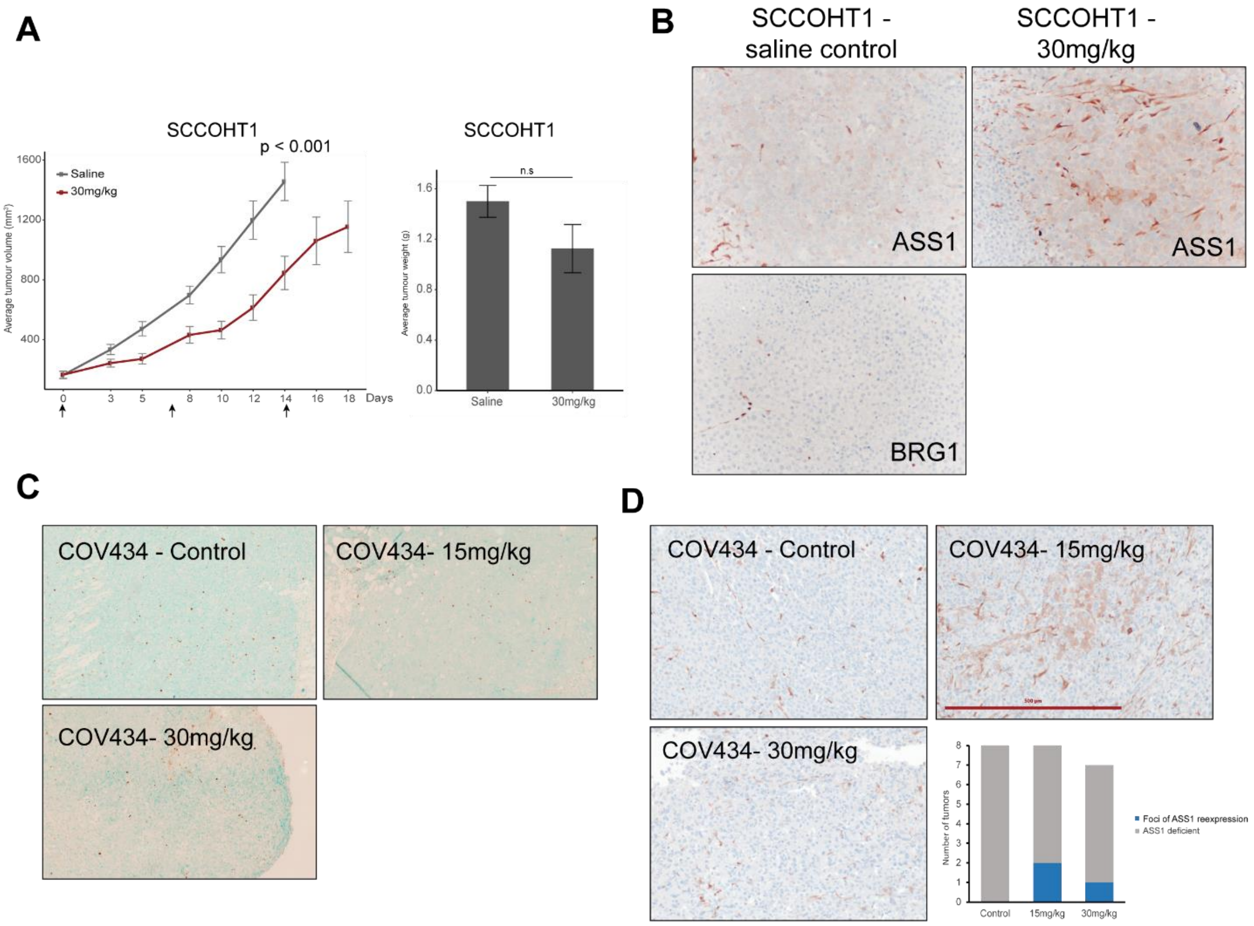


**Figure S7.** Mouse models of SCCOHT cell lines. **A,** tumor volume and weight of SCCOHT1 mouse model. Significance was calculated using ANOVA, followed by a post-hoc Tukey's test. Error bars represent standard error of mean. **B,** ASS1 re-expression in SCCOHT1 cells in both saline control group and 30mg/kg treated groups. **C,** Representative tunnel assay images from each treatment group indicating no significant increase in apoptosis in tumors from the ADI-PEG20 treated compared to saline-treated groups, and **D,** COV434 subcutaneous xenograft showing foci of ASS1 re-expression in treated groups.
